# Quantifying vector diversion effects in zoonotic systems: A modelling framework for arbovirus transmission between reservoir and dead-end hosts

**DOI:** 10.1101/2025.07.24.666561

**Authors:** Emma L Fairbanks, Matthew Baylis, Janet M Daly, Michael J Tildesley

## Abstract

Vector-borne disease transmission involves complex interactions between vectors, reservoir hosts and dead-end hosts. We present a mathematical model for the vectorial capacity that incorporates multiple host types and their interactions, focusing specifically on West Nile virus transmission by *Culex pipiens* mosquitoes. Our model integrates climate-dependent parameters affecting vector biology with vector control interventions to predict transmission potential under various scenarios. We demonstrate how vector control interventions targeting one host type can significantly impact transmission dynamics across all host populations. By examining the effects of different vector control tool modes of action (repellency, preprandial killing, disarming and postprandial killing), we develop target product profiles that minimise unintended consequences of vector control. Notably, we identify the optimal intervention characteristics needed to prevent repellency on dead-end hosts from inadvertently increasing transmission among reservoir hosts. This research provides valuable insights for public health officials designing targeted vector control strategies and offers a flexible modelling framework that can be adapted to other vector-borne diseases with complex host dynamics.

**Author summary:** Mosquitoes that spread diseases like West Nile virus don’t just bite one type of animal—they feed on birds, humans, and other mammals. This creates a complex web of disease transmission that current prevention strategies often overlook. We developed a mathematical model to understand what happens when mosquito control methods target different types of hosts in this network.

Our research reveals a surprising and concerning finding: when people use repellents to protect themselves from mosquitoes, those mosquitoes don’t simply disappear—they redirect to birds instead. Since birds are the main animals that can spread West Nile virus to other mosquitoes, this redirection can actually increase disease transmission in the bird population by up to 23%. More infected birds ultimately means more infected mosquitoes and higher risk for humans.

However, we also identified a solution. We found that if repellent products could kill just 2% of the mosquitoes they encounter before those mosquitoes find alternative hosts, this would eliminate the harmful redirection effect. Our work provides specific guidelines for developing better mosquito control products and helps public health officials understand the broader consequences of different intervention strategies. This framework can be applied to other mosquito-borne diseases beyond West Nile virus.

## 1 Introduction

While many mathematical models for predicting the risk of disease transmission consider the effects of climate and vector control on the number of vectors [1–15], sometimes the vectorial capacity is not considered. Vectorial capacity, a measure of the potential of a vector population to transmit a pathogen, is defined as the total number of potentially infectious bites that would arise from all vectors biting a single infectious host on a single day [16]. This not only considers the number of vectors, but also their ability to transmit a pathogen.

Therefore, vectorial capacity is the product of both the number of vectors and the relative vector capacity (rVC), defined as the total number of potentially infectious bites that would arise from a single vector biting a single host on a single day. The basic reproductive number, the expected number of cases to arise from a single infectious case in a susceptible population, can be expressed as the product of the vectorial capacity and duration of the host’s infectious period [17].

We previously modelled the rVC for viruses transmitted by *Culicoides* [18], considering how climatic variables affect the rate at which vectors feed, expected lifespan of vectors and the extrinsic incubation period (EIP), defined as the time required for a vector to become infectious after consuming an infected blood meal. The model also considers the effects of vector control interventions at different stages during the mosquito feeding cycle. Additionally, the model considers host selection, which is influenced by the availability of different host types and the vector’s preferences for these hosts.

However, there are some limitations to this model. For example, the model only considers one host species. While this may be informative in localised populations with only one competent host present, it is not possible to consider interactions between host species. In some cases, a disease may be mainly transmitted by a host which shows less severe clinical signs than other hosts or dead-end hosts (hosts which can be infected but cannot infect). Hosts which can both be infected and transmit a pathogen are often referred to as reservoir hosts. These hosts can amplify disease transmission, whereas infection of a dead-end host has no downstream effects of transmission. In these cases it is important to consider the interactions between hosts.

Vector control interventions targeting one host type can significantly impact transmission dynamics across all host populations. When control measures such as repellents or insecticides are applied to one host type, mosquito vectors may be diverted to alternative hosts, potentially altering disease transmission patterns.

In this study, we extend this model to consider the rVC for transmission of pathogens between host species. Here, we include reservoir hosts and dead-end hosts. As an example, we model West Nile virus (WNV) transmission by *Culex pipiens*, where interactions between reservoir hosts (birds) and dead-end hosts (humans and other mammals) determine overall transmission intensity.

## 2 Methods

### 2.1 Model development

The baseline model for simulating the dynamics in a single host species is described in full detail in Fairbanks et al. [18]. Here, the rVC on day *t* was calculated as the probability the vector feeds on the host of interest on day *t* and survives until the end of that day multiplied by the probability it infects another of the same host type every day after, given it survives until that day. The model considered the climatic dependence of the rates of gonotrophic cycle completion, vector mortality and pathogen EIP completion. It can either be simulated using daily climatic data or, alternatively, for a constant temperature or set durations of gonotrophic cycle completion, vector lifespan and EIP completion.

In this study, we extend this model to account for the interactions between vectors and different host types. This multiple host model is described in detail in the Supplementary information. It can be utilised to calculate the transmission potential from each reservoir host species to each other host species, including dead-end hosts. For interpretability, we define the basic reproduction number (*R*_0_) as the product of the rVC, the infectious period and the expected number of vectors per host [17, 18].

Within the modelling framework, vectors that encounter hosts can experience several endpoints: they might successfully feed, be killed before (preprandial mortality), become disarmed (prolonged blood feeding inhibition) or abandon the current host to search for another. Vectors that succeed in feeding may still be killed after the blood meal (postprandial mortality).

The model simulations and subsequent data processing were conducted in R, with visualisations generated using the ggplot2 package [19] within the RStudio integrated development environment [20].

### 2.2 Model application

We simulate the model for West Nile virus transmission by *Cx. pipiens*. The model parameters were set following a comprehensive literature review. The rate of gonotrophic cycle completion and vector life-span p day were parameterised by Shocket et al. [21] using data from four [22–25] and three [24, 26, 27] *Cx. pipiens* studies, respectively. For the EIP, we use the model from Vollans et al. [28], which was parameterised using Bayesian methods and data from 2145 mosquitoes. This parameterisation includes the probability of transmission from host to vector and therefore we set *ρ*_*h*→*v*_ = 1 for reservoir hosts. Vollans et al. [28] found that the gamma and Weibull distributions performed similarly. Here, we favour the gamma distribution since it has a property of being able to be broken up into multiple exponential distributions. This allows for consideration of daily fluctuations in temperature.

When estimating the number of vectors per host required for *R*_0_ > 1, we consider avian reservoir hosts with infectious periods of 7 and 10 days [29]. An infectious period of 7 days may be applicable to geese [30], passerines [31] and owls [32], and 10 days is applicable to raptors [33] and turkeys [34].

#### 2.2.1 Comparing the transmission potential between host types

Griep et al. [35] reviewed blood meal sources from 26,857 *Culex* species, including 2062 *Cx. pipiens* in palearctic regions. They found that 68.3% percent had fed on avian hosts, and 14.1% and 17.2% fed on human and non-human mammal hosts, respectively. We consider the rVC from birds (the reservoir hosts) to human and other mammal hosts (assuming these are a dead-end hosts). As not all mammals are hosts of WNV this is likely to represent an upper bound. We calculate the rVC for temperatures between 10 and 32 °C using these fixed blood selection values, however these are likely to vary between settings [36].

The probability of transmission from vector to host is assumed to be the same for all hosts, giving *ρ*_*h→v*_ = 0.74 for all three hosts [37]. We also assume that vectors bite only once per feeding cycle and that there are no vector control tools present.

#### 2.2.2 Simulation at a larger spatial scale

We then simulate the model considering daily temperature time series from the UK to produce maps showing the patterns in transmission potential. Temperature for 2019–2023 was extracted from the HadUK-Grid gridded average climate observations for 5 km grid squares of the UK [38]. For each year, we calculate the rVC each day. To summarise the results, we pool estimates for each month across days and year to calculate the monthly minimum, mean and maximum. The UK represents an ideal case study for demonstrating applicability of the model, as it encompasses considerable climatic variation across a relatively compact geographical area, allowing for examination of diverse transmission dynamics.

#### 2.2.3 Considering the effect of vector-control tools on different host types

When considering how vector control affects the transmission potential, we will consider two host types; a reservoir (birds) and another dead-end host. We vary the percentage of vectors feeding on the reservoir host and assume the rest feed on a dead-end host. We simulate the model for settings where, in the absence of vector control, the vector would select a reservoir host 20%, 50% or 80% of the time to capture a range of demographic settings.

Initially, we will consider the effects of a tool which only repels mosquitoes, therefore they continue host-seeking and may bite another host of any type. We vary the repellent effect assuming a 0–100% reduction in the rate of blood feeding. The coverage is also varied from 0-100% of the target host type having the tool. We assume that if a host type has access to a tool they always use it when exposed to mosquitoes. When analysing the effects of vector-control we fix the temperature at the optimal value for transmission, as calculated in Section 2.2.1.

Finally, we investigate how other vector control characteristics (modes of action) can counteract the potential negative effects of repellents. When repellents are applied to dead-end hosts, they may inadvertently increase transmission by redirecting mosquitoes towards reservoir hosts. For each percentage of the reduction in the rate of biting investigated (0-100%), we therefore calculate the percentage of the repelled mosquitoes which would need to follow each alternative mode of action (preprandial killing, disarming, or postprandial killing) to mitigate this increased transmission risk.

## 3 Results

### 3.1 Model development

Our study extends a mathematical model of the rVC that integrates climatic factors and vector control interventions, introducing interactions between vectors and multiple types of host. This approach enables consideration of secondary and dead-end hosts in the analysis allowing for consideration of how control tools on one host type affect transmission potential to other host types, as demonstrated in the following results.

### 3.2 Comparing the transmission potential to host types

As expected, the magnitude of the rVC for each type of host corresponds to the likelihood a vector selects the host, with hosts fed upon more often receiving a larger portion of the disease burden (Figure 1). The maximum rVC is when the temperature is 24.5 °C. This corresponds to a rVC of 0.061, 0.013 and 0.015 for birds, humans and other mammals, respectively.

**Figure 1.**
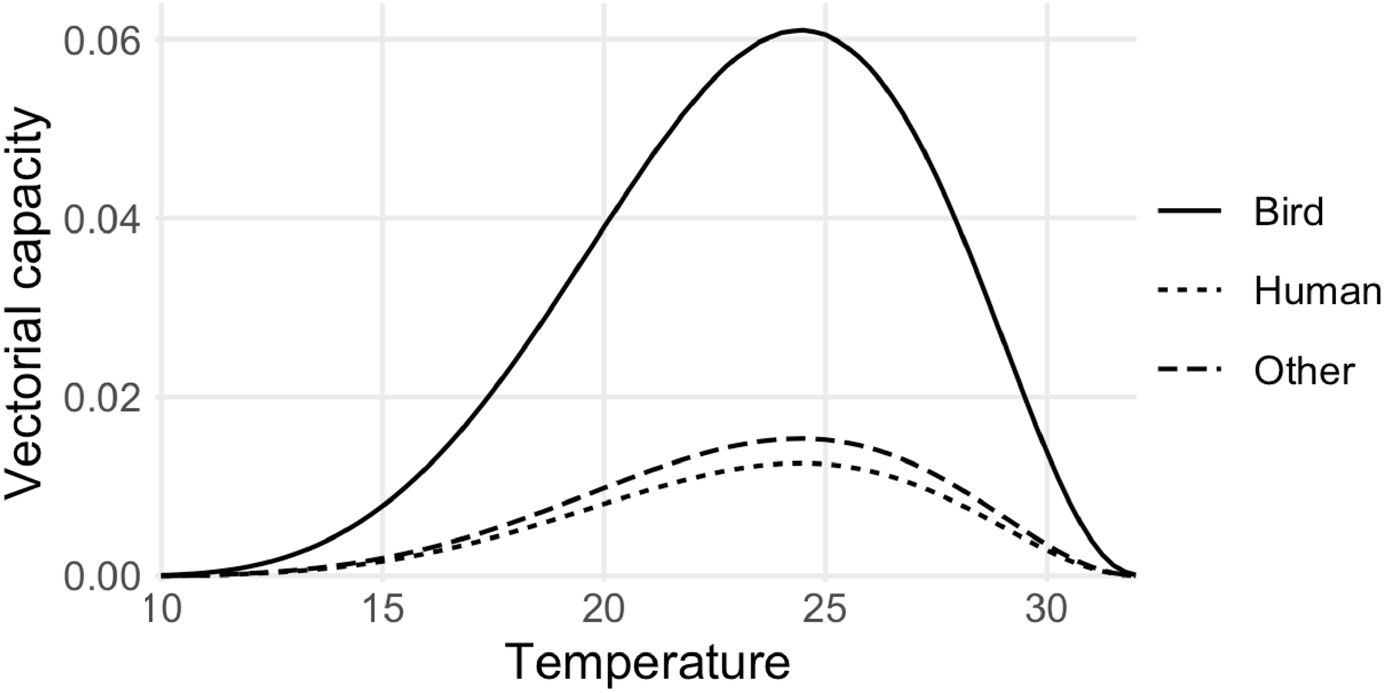
The relative vectorial capacity from birds (the reservoir host) to each host type at constant temperatures.

For a bird species with an infectious period of 7 days, the vectors per host required for *R*_0_ > 1 in birds, humans and other hosts would be 2.34, 10.99 and 9.52, respectively. For bird species with an infectious period of 10 days the required number of vectors per host would be 1.64, 7.69 and 6.67 for birds, humans and other hosts, respectively.

### 3.3 Large scale simulation

Using daily mean temperature data, we produced maps of the mean monthly rVC (Figure 2). This allows for comparisons both spatially and temporally. Further details regarding monthly rVC values and corresponding vector population thresholds can be found in Supplementary table S3.

**Figure 2.**
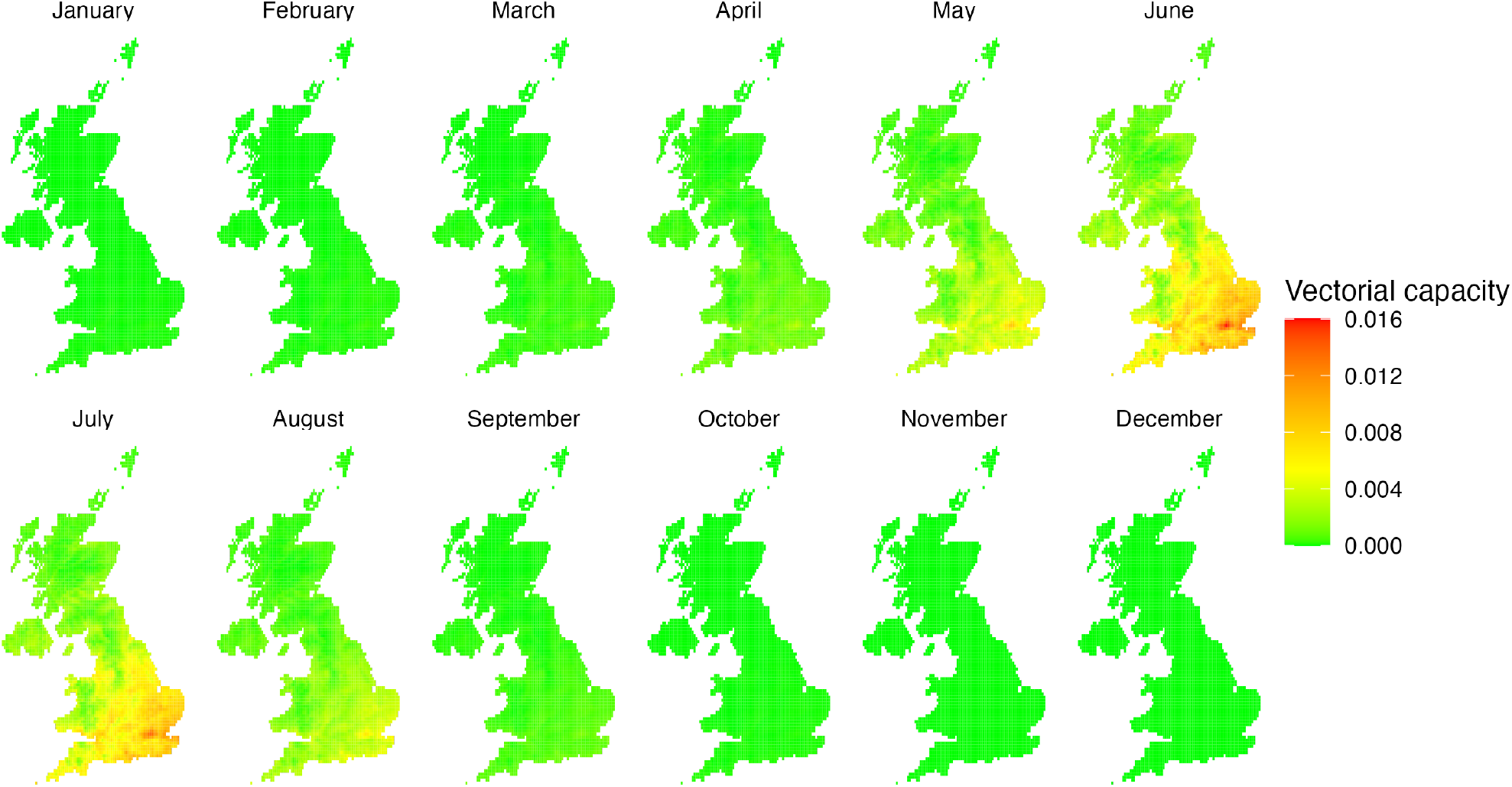
Monthly mean predicted relative vectorial capacity for birds (the reservoir host) calculated for mean daily temperature estimates for 2019–2023.

The modelling outputs reveal considerable spatial heterogeneity. We observe that the rVC in Scotland and Northern Ireland is relatively low when compared with Wales and England, with England having the largest predicted rVC on average. The model predicts the largest rVC in south-east England, however, similar values are observed across the south of England, in South Wales and as far north as Merseyside and parts of Yorkshire.

The estimates demonstrate pronounced seasonal variations in rVC with significant implications for disease transmission dynamics. June exhibits the peak rVC (0.016), situated within a broader high-transmission window extending from May to August. During this summer period, the required vector population threshold for disease establishment (*R*_0_ > 1) remains biologically plausible, ranging from approximately 6 to 15 vectors per host for an avian reservoir host with a 10-day infectious period.

In contrast, the winter months (November to March) show markedly reduced transmission potential, with numerous locations experiencing a rVC of zero. The maximum rVC in December is approximately 1500 times lower than the June peak, corresponding to an extraordinarily high vector population threshold (9371.54 vectors per host) that renders sustained transmission biologically unlikely given typical winter vector population densities. April and September-October emerge as critical transition periods, characterised by rapidly changing rVC as environmental conditions shift.

### 3.4 Considering the effect of vector-control tools on different host types

For interpretation, it is important to consider that, given a fixed number of vectors per host within a population, we would expect the vectorial capacity to increase or decrease proportionally with the rVC. This assumes that the size of the vector population is limited by the carrying capacity, which is a common assumption when modelling the impact of interventions on vectorial capacity [39–41].

Figure 3 shows that while repelling vectors from a reservoir host always reduces the rVC of the reservoir and dead-end hosts, repelling vectors from dead-end hosts can increase the rVC of reservoir hosts. As coverage of the tool and repellency increase we see a larger decrease in rVC for both host types when the tool is given to a reservoir host or in dead-end hosts when the tool is given to a dead-end host, and a larger increase in the rVC in a reservoir host when the tool is given to a dead-end host.

**Figure 3.**
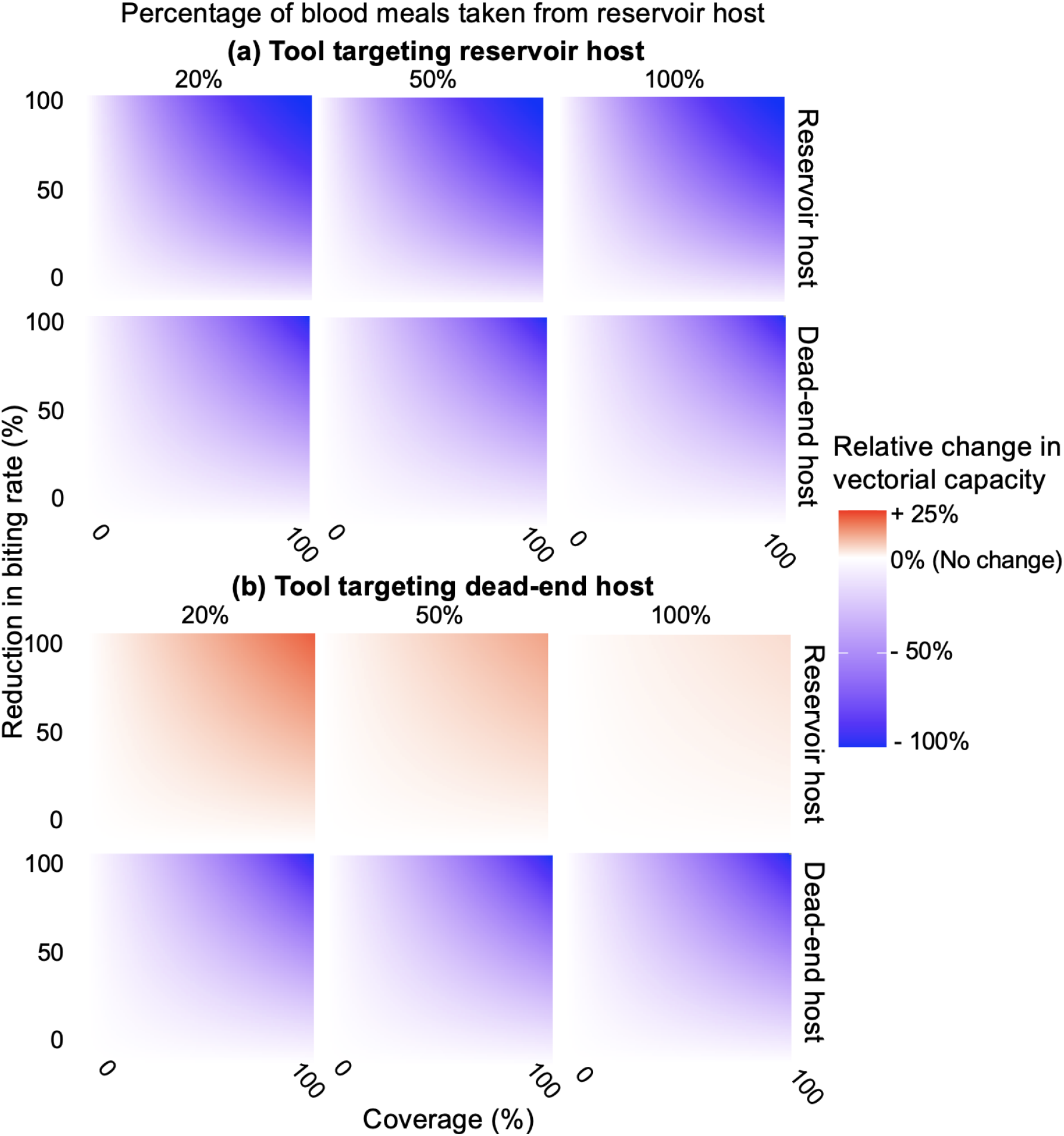
The estimated relative change in vectorial capacity for the reservoir and dead-end host when a tool is used by a (a) reservoir or (b) dead-end host for tools which reduce the rate of biting at different population coverage levels.

When targeting dead-end hosts, the reduction in the rVC of dead-end hosts tells us that each mosquito that feeds on an infectious reservoir is less likely to cause an infection in the dead-end host. However, the unintentional increase in vectorial capacity in reservoir hosts suggests that there will be more infected reservoir hosts, and therefore potentially more vectors becoming infected. In our scenario, where the model is parameterised to describe WNV transmission by *Cx. pipiens*, we see an increase in vectorial capacity of up to 23.4%.

### 3.5 Target product profiles for vector-control tools

For a given level of repellency for a tool used on a dead-end host, Figure 4 shows the estimated magnitude of the other modes of action required to avoid the increase in vectorial capacity for reservoir hosts. Tools which preprandially kill at least 2% of *Cx. pipiens* of the repelled vectors counteract these unintended negative effects for all levels of repellency. However, for tools which repel almost all (> 96%) mosquitoes feeding on dead-end hosts, postprandially killing or disarming cannot counteract all of the negative effects of the repellency.

**Figure 4.**
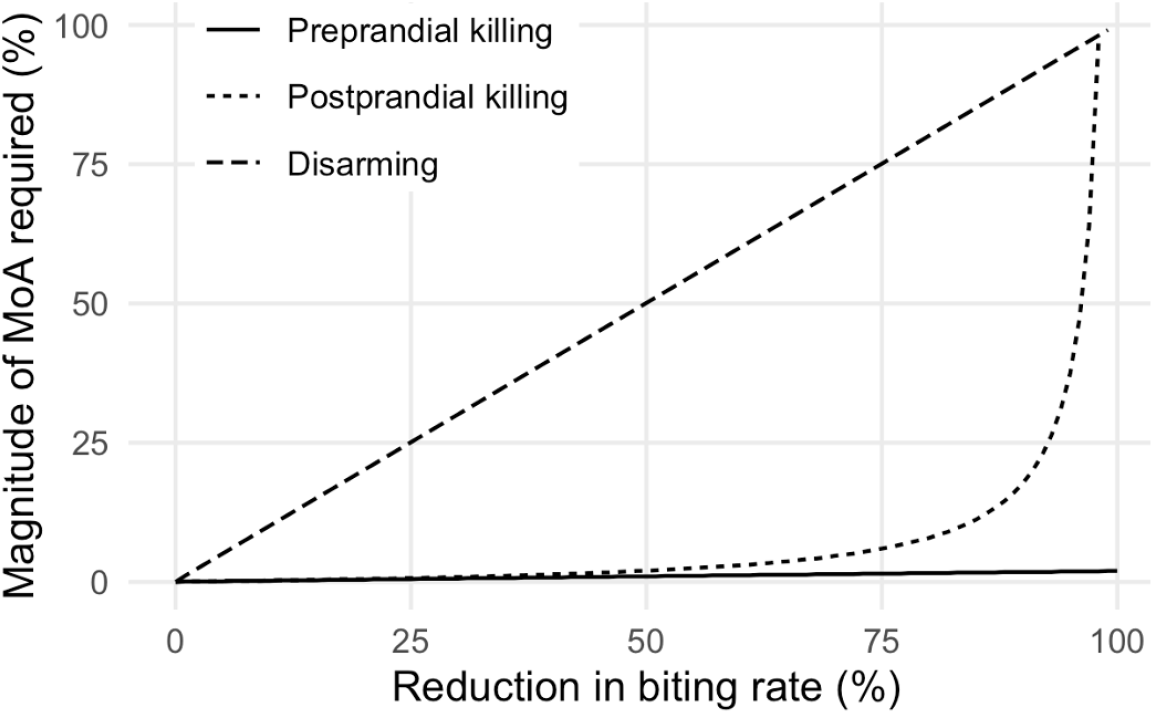
Magnitude of each mode of action (MoA) required to counteract the increased vectorial capacity in reservoir hosts when a tool repells hosts from dead-end hosts.

Postprandial mortality is more effective than disarming, however its effects rapidly decrease when over 80% of vectors are repelled. The percentage of mosquitoes which need to be postprandially killed, given 80% or 90% of mosquitoes are repelled, is 8% or 18%, further increasing to 38% when 95% of mosquitoes are repelled. This is because the vectors must feed for postprandial killing to be effective – the fewer vectors that feed, of those that do feed a higher percentage need to be killed.

## 4 Discussion

To our knowledge, this is the first model that quantifies the rVC between multiple host types independently. Understanding these complex interactions is crucial for developing effective vector control strategies, as interventions focused solely on protecting dead-end hosts could inadvertently increase transmission among reservoir hosts, potentially elevating overall disease risk. By modelling these relationships, we can identify optimal intervention characteristics that minimise unintended consequences of vector control and maximise public health benefits.

The optimal temperature for WNV transmission was estimated to be 24.5°C, in line with previous estimates [21, 42]. This temperature represents the thermal optimum where rVC reaches its peak due to the combined effects of accelerated mosquito development, shortened gonotrophic cycle, and reduced EIP, while still maintaining sufficient vector longevity. Furthermore, this optimal temperature falls within the summer temperature range of many temperate regions, including parts of the UK, explaining the seasonal patterns observed in our spatial analysis. For the UK, modelled seasonal and spatial patterns in rVC suggest that early warning systems, disease surveillance and control strategies would benefit from targeted implementation, with intensified efforts during the high-transmission window and in locations exhibiting elevated rVC.

Our model predicts higher rVC values in urban areas, particularly London. Urban areas typically experience temperatures 1–3°C higher than surrounding rural environments [43]. This urban amplification can be attributed to the urban heat island effect, which creates microclimatic conditions that are more favourable for *Cx. pipiens*, potentially extending the transmission season and intensifying WNV risk in temperate, metropolitan areas. Additionally, recent research indicates that while *Cx. pipiens* larvae are more abundant in urban residential areas due to the prevalence of artificial breeding habitats such as storm drains, adult mosquitoes often show higher densities in urban parks where they find more suitable microclimatic conditions and diverse hosts [44]. While our model focuses primarily on rVC, it is important to acknowledge that actual disease transmission risk is the product of both rVC and the number of vectors per host. Mosquito population dynamics are influenced by numerous factors beyond temperature, including rainfall patterns and habitat availability. The heterogeneity in vector abundance can significantly modulate transmission risk even in areas with high rVC.

Host selection of vectors depends on host preference and host availability [18], therefore is likely to vary across settings. Experimental studies have demonstrated that *Cx. pipiens* prefer some avian species over others [45]. However, these preferences are modulated by environmental conditions and host availability in field settings, leading to spatial and temporal variation in feeding patterns [**?**]. Such heterogeneity in host selection across different landscapes significantly impacts WNV transmission potential.

This modelling framework is flexible for application to other vectors and pathogens. Due to a reduction in wildlife habitats, we are observing an increase in the risk of human exposure to emerging and existing zoonotic pathogens [46]. An increase in transmission between wildlife reservoirs and human hosts may also have implications for disease elimination. Currently, numbers of human zoonotic malaria cases are increasing globally and there are currently no control measures that target wildlife reservoirs [46, 47].

Many health agencies suggest using repellent to reduce the risk of vector-borne disease [48– 50]]. This work shows that for zoonotic diseases this may not always be the case. More investigation is needed into the effects of increasing vectorial capacity on disease transmission. The modes of action of vector control tools change according to the resistance level of vectors and insecticide dose [39, 40]. Insecticide resistance has been reported in *Cx. pipiens* in Europe [51] and globally [52]. Therefore, more data on the responses of local vector populations to insecticides should be gathered to ensure robust evaluation within different settings. This modelling framework can be used to assess whether these tools are suitable for use on dead-end hosts, and suggest target product profiles of additional tools for use.

Repellents that also kill or disarm mosquitoes are more likely to avoid unintentional increases in vectorial capacity in reservoir hosts, with preprandial killing being the most effective. This study provides the first quantitative estimates of the magnitude of unique modes of action of tools required to reduce disease transmission. These quantitative thresholds offer valuable targets for product developers and public health officials designing next-generation vector control strategies. In our WNV example, if 2% of the reduction in biting is due to preprandial killing, vectorial capacity will be reduced in all hosts. However, this may vary across different settings and vector populations [41].

Our mathematical modelling presented here advances the understanding of host-vector-pathogen interactions. This work demonstrates the critical importance of considering entire transmission systems when implementing vector control, as interventions targeting one host population can significantly influence transmission throughout the ecological network. This underscores the need for integrated approaches to vector control that consider both personal protection and community-level impacts. The model is a valuable tool for assessing transmission risk and optimising control interventions in the face of emerging zoonotic threats and changing climate conditions.

## Conflict of interest

The authors declare no conflict of interest.

## Ethical approval

The authors confirm that the ethical policies of the journal, as noted on the journal’s author guidelines page, have been adhered to. No ethical approval was required.

## Author contributions

### Emma L Fairbanks

Conceptualisation, Methodology, Software, Validation, Formal analysis, Investigation, Data Curation, Writing - Original Draft, Writing - Review & Editing, Visualisation, Funding acquisition; **Matthew Baylis:** Conceptualisation, Writing - Review & Editing; **Janet M Daly:** Conceptualisation, Writing - Review & Editing, Funding acquisition; **Michael J Tildesley:** Conceptualisation, Methodology, Writing - Review & Editing, Funding acquisition

### Financial support

The authors were supported by the Horserace Betting Levy Board (vet/prj/809). MJT is funded on a joint BBSRC/EEID grant (BB/T004312/1).

### Data Availability Statement

Code to simulate the model will be made available in a GitHub package and achieved using Zenodo to provide a DOI upon publication. This is provided for review in VectorialCapacity_MultipleHosts.R.

## Notes

### Competing Interest Statement

The authors have declared no competing interest.

## References

[1] AJ Worton, RA Norman, L Gilbert, and RB Porter. GIS-ODE: linking dynamic population models with GIS to predict pathogen vector abundance across a country under climate change scenarios. J R Soc Interface, 21(217):20240004, 2024. doi: 10.1098/rsif.2024.0004.

[2] LHV Franklinos, DW Redding, TCD Lucas, R Gibb, I Abubakar, and KE Jones. Joint spatiotemporal modelling reveals seasonally dynamic patterns of Japanese encephalitis vector abundance across India. PLoS Negl Trop Dis, 16(2):e0010218, 2022. doi: 10.1371/journal.pntd.0010218.

[3] Modelling the monthly abundance of Culicoides biting midges in nine european countries using random forests machine learning. Parasit Vectors, 13(1):194, 2020. doi: 10.1186/s13071-020-04053-x.

[4] J Liu-Helmersson, A Brännström, MO Sewe, JC Semenza, and J Rocklöv. Estimating. past, present, and future trends in the global distribution and abundance of the arbovirus vector Aedes aegypti under climate change scenarios. Front Public Health, 7: 148, 2019. doi: 10.3389/fpubh.2019.00148.

[5] MC Cecere, LI Rodríguez-Planes, GM Vazquez-Prokopec, U Kitron, and RE Gürtler. Community-based surveillance and control of chagas disease vectors in remote rural areas of the Argentine Chaco: A five-year follow-up. Acta Trop, 191:108–115, 2019. doi: 10.1016/j.actatropica.2018.12.038.

[6] RS McCann, JP Messina, DW MacFarlane, MN Bayoh, JE Gimnig, E Giorgi, and ED Walker. Explaining variation in adult Anopheles indoor resting abundance: the relative effects of larval habitat proximity and insecticide-treated bed net use. Malar J, 16(1):288, 2017. doi: 10.1186/s12936-017-1938-1.

[7] HE Brown, R Barrera, AC Comrie, and J Lega. Effect of temperature thresholds on modeled Aedes aegypti (Diptera: Culicidae) population dynamics. J Med Entomol, 54 (4):869–877, 2017.

[8] DA Ewing, CA Cobbold, BV Purse, MA Nunn, and SM White. Modelling the effect of temperature on the seasonal population dynamics of temperate mosquitoes. J Theor Biol, 400:65–79, 2016. doi: 10.1016/j.jtbi.2016.04.008.

[9] G Marini, P Poletti, M Giacobini, A Pugliese, S Merler, and R Rosà. The role of climatic and density dependent factors in shaping mosquito population dynamics: the case of Culex pipiens in northwestern Italy. PLoS One, 11(4):e0154018, 2016. doi: 10.1371/journal.pone.0154018.

[10] C Christiansen-Jucht, K Erguler, CY Shek, M-G Basáñez, and PE Parham. Modelling Anopheles gambiae ss population dynamics with temperature-and age-dependent survival. Int J Environ Res Public Health, 12(6):5975–6005, 2015. doi: 10.3390/ijerph120605975.

[11] C Talla, D Diallo, I Dia, Y Ba, J-A Ndione, AA Sall, A Morse, A Diop, and M Diallo. Statistical modeling of the abundance of vectors of West african Rift Valley fever in Barkédji, Senegal. PLoS One, 9(12):e114047, 2014. doi: 10.1371/journal.pone.0114047.

[12] PE Parham, D Pople, C Christiansen-Jucht, S Lindsay, W Hinsley, and E Michael. Modeling the role of environmental variables on the population dynamics of the malaria vector Anopheles gambiae sensu stricto. Malar J, 11:271, 2012. doi: 10.1186/1475-2875-11-271.

[13] J Wang, NH Ogden, and H Zhu. The impact of weather conditions on Culex pipiens and Culex restuans (Diptera: Culicidae) abundance: a case study in Peel region. J Med Entomol, 48(2):468–475, 2011. doi: 10.1603/me10117.

[14] KP Paaijmans, SS Imbahale, MB Thomas, and W Takken. Relevant microclimate for determining the development rate of malaria mosquitoes and possible implications of climate change. Malar J, 9:196, 2010. doi: 10.1186/1475-2875-9-196.

[15] B Schaeffer, B Mondet, and S Touzeau. Using a climate-dependent model to predict mosquito abundance: application to Aedes (Stegomyia) africanus and Aedes (Diceromyia) furcifer (Diptera: Culicidae). Infect Genet Evol, 8(4):422–432, 2008. doi: 10.1016/j.meegid.2007.07.002.

[16] C Garrett-Jones. The human blood index of malaria vectors in relation to epidemiological assessment. Bull World Health Organ, 30(2):241, 1964.

[17] SPC Brand and MJ Keeling. The impact of temperature changes on vector-borne disease transmission: Culicoides midges and bluetongue virus. J R Soc Interface, 14 (128):20160481, 2017. doi: 10.1098/rsif.2016.0481.

[18] EL Fairbanks, JM Daly, and MJ Tildesley. Modelling the influence of climate and vector control interventions on arbovirus transmission. Viruses, 16(8):1221, 2024. doi: 10.3390/v16081221.

[19] H Wickham, W Chang, L Henry, TL Pedersen, K Takahashi, C Wilke, K Woo, H Yutani, D Dunnington, and RStudio. Create Elegant Data Visualisations Using the Grammar of Graphics, 2022. URL https://ggplot2.tidyverse.org.

[20] RStudio Team. Rstudio: Integrated Development Environment for R. RStudio, PBC., Boston, MA, 2020. URL http://www.rstudio.com/.

[21] Marta S Shocket, Anna B Verwillow, Mailo G Numazu, Hani Slamani, Jeremy M Cohen, Fadoua El Moustaid, Jason Rohr, Leah R Johnson, and Erin A Mordecai. Transmission of West Nile and five other temperate mosquito-borne viruses peaks at temperatures between 23 c and 26 c. Elife, 9:e58511, 2020. doi: 10.7554/eLife.58511.

[22] J Li, G Zhu, H Zhou, J Tang, and J Cao. Effect of temperature on the development of Culex pipiens pallens. Chin J Vector Biol Control, 28:35–37, 2017.

[23] DJ Madder, GA Surgeoner, and BV Helson. Number of generations, egg production, and developmental time of Culex pipiens and Culex restuans (Diptera: Culicidae) in southern Ontario. J Med Entomol, 20(3):275–287, 1983. doi: 10.1093/jmedent/20.3.275.

[24] Jordan E Ruybal, Laura D Kramer, and AM Kilpatrick. Geographic variation in the response of Culex pipiens life history traits to temperature. Parasit Vectors, 9:116, 2016. doi: 10.1186/s13071-016-1402-z.

[25] A Tekle. The physiology of hibernation and its role in the geographical distribution of populations of the Culex pipiens complex. Am J Trop Med Hyg, 9:321–330, 1960. doi: 10.4269/ajtmh.1960.9.321.

[26] SS Andreadis, OC Dimotsiou, and M Savopoulou-Soultani. Variation in adult longevity of Culex pipiens f. pipiens, vector of the West Nile virus. Parasitol Res, 113:4315–4319, 2014. doi: 10.1007/s00436-014-4152-x.

[27] AT Ciota, AC Matacchiero, AM Kilpatrick, and LD Kramer. The effect of temperature on life history traits of Culex mosquitoes. J Med Entomol, 51(1):55–62, 2014. doi: 10.1603/me13003.

[28] M Vollans, J Day, S Cant, J Hood, AM Kilpatrick, LD Kramer, A Vaux, J Medlock, T Ward, and RS Paton. Modelling the temperature dependent extrinsic incubation period of West Nile virus using Bayesian time delay models. J Infect, 89(6):106296, 2024. doi: 10.1016/j.jinf.2024.106296.

[29] V Gamino and U Höfle. Pathology and tissue tropism of natural West Nile virus infection in birds: a review. Vet Res, 44(1):39, 2013. doi: 10.1186/1297-9716-44-39.

[30] C Banet-Noach, L Simanov, and M Malkinson. Direct (non-vector) transmission of West Nile virus in geese. Avian Pathol, 32(5):489–494, 2003. doi: 10.1080/0307945031000154080.

[31] DA LaPointe, EK Hofmeister, CT Atkinson, RE Porter, and RJ Dusek. Experimental infection of hawaiiamakihi (Hemignathus virens) with West Nile virus and competence of a co-occurring. vector, Culex quinquefasciatus: Potential impacts on endemic hawaiian avifauna. J Wildl Dis, 45(2):257–271, 2009. doi: 10.7589/0090-3558-45.2.257.

[32] NM Nemeth, DC Hahn, DH Gould, and RA Bowen. Experimental West Nile virus infection in eastern screech owls (Megascops asio). Avian Dis, 50(2):252–258, 2006. doi: 10.1637/7466-110105R1.1.

[33] U Ziegler, J Angenvoort, D Fischer, C Fast, M Eiden, AV Rodriguez, S Revilla-Fernández, N Nowotny, JG de la Fuente, M Lierz, and MH Groschup. Pathogenesis of West Nile virus lineage 1 and 2 in experimentally infected large falcons. Vet Microbiol, 161(3-4):263–273, 2013. doi: 10.1016/j.vetmic.2012.07.041.

[34] DE Swayne, JR Beck, and S Zaki. Pathogenicity of West Nile virus for turkeys. Avian Dis, pages 932–937, 2000. doi: 10.2307/1593067.

[35] JS Griep, E Grant, J Pilgrim, O Riabinina, M Baylis, M Wardeh, and MSC Blagrove. Meta-analyses of Culex blood-meals indicates strong regional effect on feeding patterns. PLoS Negl Trop Dis, 19(1):e0012245, 2025. doi: 10.1371/journal.pntd.0012245.

[36] Emma L Fairbanks, Michael J Tildesley, and Janet M Daly. A systematic review quantifying host feeding patterns of culicoides species responsible for pathogen transmission. bioRxiv, pages 2024–07, 2024. doi: 10.1101/2024.07.25.605155.

[37] Marjorie J Wonham, Mark A Lewis, Joanna Rencławowicz, and P Van den Driessche. Transmission assumptions generate conflicting predictions in host–vector disease models: a case study in West Nile virus. Ecol Lett, 9(6):706–725, 2006. doi: 10.1111/j.1461-0248.2006.00912.x.

[38] D Hollis, M McCarthy, M Kendon, T Legg, and I Simpson. HadUK-grid‚ÄîA new UK dataset of gridded climate observations. Geosci Data J, 6(2):151–159, 2019. doi: 10.1002/gdj3.78.

[39] EL Fairbanks, M Saeung, A Pongsiri, E Vajda, Y Wang, DJ McIver, JH Richardson, A Tatarsky, NF Lobo, SJ Moore, A Ponlawat, T Chareonviriyaphap, A Ross, and N Chitnis. Inference for entomological semi-field experiments: Fitting a mathematical model assessing personal and community protection of vector-control interventions. Comput Biol Med, 168:107716, 2024. doi: 10.1016/j.compbiomed.2023.107716.

[40] EL Fairbanks, MM Tambwe, J Moore, A Mpelepele, NF Lobo, R Mashauri, N Chitnis, and SJ Moore. Evaluating human landing catches as a measure of mosquito biting and the importance of considering additional modes of action. Sci Rep, 14(1):11476, 2024. doi: 10.1038/s41598-024-61116-0.

[41] Y Wang, N Chitnis, and EL Fairbanks. Optimizing malaria vector control in the Greater Mekong Subregion: a systematic review and mathematical modelling study to identify desirable intervention characteristics. Parasit Vectors, 17(1):162, 2024. doi: 10.1186/s13071-024-06234-4.

[42] J Heidecke, J Wallin, P Fransson, P Singh, H Sjödin, PC Stiles, M Treskova, and J Rocklöv. Uncovering temperature sensitivity of West Nile virus transmission: Novel computational approaches to mosquito-pathogen trait responses. PLoS Comput Biol, 21(3):e1012866, 2025. doi: 10.1371/journal.pcbi.1012866.

[43] NB Grimm, SH Faeth, NE Golubiewski, CL Redman, J Wu, X Bai, and JM Briggs. Global change and the ecology of cities. Science, 319(5864):756–760, 2008. doi: 10.1126/science.1150195.

[44] L Krol, M Langezaal, L Budidarma, D Wassenaar, EA Didaskalou, K Trimbos, M Dellar, PM van Bodegom, GW Geerling, and M Schrama. Distribution of Culex pipiens life stages across urban green and grey spaces in. Leiden, The Netherlands. Parasit Vectors, 17(1):37, 2024. doi: 10.1186/s13071-024-06120-z.

[45] JE Simpson, CM Folsom-O’Keefe, JE Childs, LE Simons, TG Andreadis, and MA Diuk-Wasser. Avian host-selection by Culex pipiens in experimental trials. PLoS One, 4(11): e7861, 2009. doi: 10.1371/journal.pone.0007861.

[46] F Keesing and RS Ostfeld. Impacts of biodiversity and biodiversity loss on zoonotic diseases. Proc Natl Acad Sci USA, 118(17):e2023540118, 2021. doi: 10.1073/pnas.2023540118.

[47] KM Fornace, CJ Drakeley, KA Lindblade, J Jelip, and K Ahmed. Zoonotic malaria requires new policy approaches to malaria elimination. Nat Commun, 14(1):5750, 2023. doi: 10.1038/s41467-023-41546-6.

[48] Centers for Disease Control and Prevention. Preventing West Nile Virus, 2024. URL https://www.cdc.gov/west-nile-virus/prevention/index.html. Accessed: 15/05/2025.

[49] Canadian Centre for Occupational Health and Safety. West Nile Virus, 2021. URL https://www.ccohs.ca/oshanswers/diseases/westnile.html. Accessed: 15/05/2025.

[50] European Centre for Disease Prevention and Control. West Nile virus season in full swing in Europe, 2024. URL https://www.ecdc.europa.eu/en/news-events/west-nile-virus-season-full-swing-europe. Accessed: 15/05/2025.

[51] S Vereecken, A Vanslembrouck, IM Kramer, and R Müller. Phenotypic insecticide resistance status of the Culex pipiens complex: a european perspective. Parasit Vectors, 15(1):423, 2022. doi: 10.1186/s13071-022-05542-x.

[52] JG Scott, MH Yoshimizu, and S Kasai. Pyrethroid resistance in Culex pipiens mosquitoes. Pestic Biochem Physiol, 120:68–76, 2015. doi: 10.1016/j.pestbp.2014.12.018.

